# Intersubject representational similarity analysis uncovers the impact of state anxiety on brain activation patterns in the human extrastriate cortex

**DOI:** 10.1101/2023.07.29.551128

**Authors:** Po-Yuan A. Hsiao, M. Justin Kim, Feng-Chun B. Chou, Pin-Hao A. Chen

## Abstract

The current study used functional magnetic resonance imaging (fMRI) and showed that state anxiety modulated extrastriate cortex activity in response to emotionally-charged visual images. State anxiety and neuroimaging data from 53 individuals were subjected to an intersubject representational similarity analysis (ISRSA), wherein the geometries between neural and behavioral data were compared. This analysis identified the extrastriate cortex (fusiform gyrus and area MT) to be the sole regions whose activity patterns covaried with state anxiety. Importantly, we show that this brain-behavior association is revealed when treating state anxiety data as a multidimensional response pattern, rather than a single composite score. This suggests that ISRSA using multivariate distances may be more sensitive in identifying the shared geometries between self-report questionnaires and brain imaging data. Overall, our findings demonstrate that a transient state of anxiety may influence how visual information – especially those relevant to the valence dimension – is processed in the extrastriate cortex.

## Introduction

Emotions constitute an inherent aspect of human nature, profoundly shaping how we navigate the world. Among the various emotions we experience on a daily basis, anxiety holds particular significance for our survival (1–4). An adequate amount of anxiety can heighten our alertness and vigilance, facilitating the detection of threats and avoidance of dangerous situations (5). Although previous research demonstrated emotional states - anxiety in particular - can impact visual perception (6, 7), the manner in which transient states of anxiety influence neural representations of the visual world remains largely unknown. Emphasis on trait anxiety in previous studies (8, 9) created a noticeable gap in the neuroimaging literature exploring the link between state anxiety and evoked neural activity during experimental tasks. Moreover, previous functional magnetic resonance imaging (fMRI) studies employing task-based experimental designs have demonstrated its suitability in capturing individual differences in mental or emotional states (10–12), but not traits (13). Therefore, the present study aims to utilize task-based fMRI, wherein participants will be presented with emotionally valenced pictures, to examine the association between state anxiety and brain activity patterns. To achieve this, intersubject representational analysis (ISRSA) will be employed as it is well-suited for comparing similarities in multivariate patterns across diverse measurements (14–16). The ISRSA begins with computing the intersubject similarity matrix across individuals in each of the two measurements, such as brain activity or self-reported questionnaire. We then examine the secondary correlation between the two intersubject similarity matrices derived from each measurement. In other words, our aim is to examine which brain regions where participants exhibit similar responses when their self-reported responses on the questionnaire are also similar. Prior research has demonstrated that compared to univariate analysis, ISRSA is more sensitive in detecting shared geometries between the brain and behavior (14, 17). As such, the present study incorporates ISRSA and a data-driven approach to investigate which brain regions might reveal significant associations between individual variations in state anxiety and neural representations of the three valence conditions (e.g., positive, negative, or neutral). Here, we hypothesize that, compared to a single aggregate score, using a multidimensional representation might be better at capturing individual variations in state anxiety. Based on prior empirical work, we hypothesize that brain regions known to be responsive to emotionally valenced stimuli such as the amygdala, ventromedial prefrontal cortex (vmPFC), dorsomedial prefrontal cortex (dmPFC) (1, 8, 9), and the fusiform gyrus (18–20) will reveal significant associations with state anxiety.

## Method

### Participants

Fifty-six participants with no history of mental or neurological disorders were recruited for this study in 2015. This study consisted of two phases, where all participants underwent fMRI scanning in the first phase, followed by the completion of a battery of psychological questionnaires in the second phase. Three out of 56 participants failed to complete psychological questionnaires, resulting in a final sample size of 53 participants (12 males, 41 females; mean age = 20.02, age std = 1.82; 13 Asian, 4 Black, 3 Hispanic, 33 White; all participants had no history of mental or neurological disorders) for the subsequent analysis. The sample size of the present study was determined through reference to a prior investigation employing a comparable experimental design, wherein Wagner and Heatherton (21) recruited 48 participants for their study. All participants provided informed consent, adhering to the guidelines established by the Committee for the Protection of Human Subjects at Dartmouth College.

### Experimental procedure

In this study, participants were instructed to complete an emotional reactivity task in the MRI scanner. Within this task, participants were instructed to judge each scene as either indoor or outdoor in a block design paradigm. Each block consisted of six scenes sharing the same valence (i.e., positive, negative, or neutral), with each scene displayed on the screen for a duration of 2.5 seconds, resulting in a total block duration of 15 seconds. Each emotional scene block was then alternated with a fixation block lasting for 15 seconds. Five positive, five neutral, and another five negative emotional blocks were presented in a counterbalanced order across participants.

### Stimuli

A total of 90 emotional scenes were selected from the International Affective Picture System (22) to be used as stimuli in the emotional reactivity task. Among them, thirty scenes were classified as negative, meeting criteria of both unpleasant and high arousal (valence M = 2.8; arousal M = 5.1). Similarly, thirty scenes were categorized as positive, reflecting pleasant and high arousal (valence M = 7.2; arousal M = 5.0). Additionally, a set of thirty scenes, characterized by neither pleasant nor unpleasant and low arousal, were classified as neutral (valence M = 5.4; arousal M = 3.6). To prevent potential confounds arising from an imbalance in the presence of faces or people across emotional conditions, we ensured that the ratio of faces or people presence was equivalent across the three emotional conditions. This step aimed to rule out any biases that could attribute observed differences in response to a particular emotional condition due to an over-abundance of faces or people, following the same procedure described previously (21).

### State anxiety

After the fMRI scanning session, participants were asked to complete the State-Trait Anxiety Inventory (STAI) (23). The STAI is a widely-used scale to assess individual differences in both their state and trait anxiety. Specifically, the State Anxiety subscale, consisting of 20 items, measures the current and transient state of anxiety stemming from stress or other negative emotions. State anxiety tends to decrease as stress is reduced or negative emotions are alleviated. In contrast, the Trait Anxiety subscale, also containing 20 items, captures enduring individual differences in the stable and long-term propensity for anxiety. Trait anxiety, as opposed to transient states, arises from inherent or long-standing stress factors, rendering it more stable and less susceptible to short-term fluctuations in external stimuli or events. Given the primary focus of this study on individual differences in state anxiety, our following reported results would focus on the findings related to state anxiety. However, in order to compare the findings of state anxiety, we also reported our findings related to trait anxiety.

### Image acquisition and preprocessing

#### Imaging parameters

MRI scanning was conducted using a Philips Achieva 3T scanner with a 32-channel head coil. Structural images were acquired by using a T1-weighted MPRAGE protocol, consisting of 160 sagittal slices with a TR of 9.9 ms, TE of 4.6 ms, voxel size of 1 × 1 x 1 mm, and flip angles of 8 degrees). Functional images were acquired using a T2^*∗*^-weighted echo-planar sequence, consisting of 36 axial slices, 3 mm thick with 0.5 mm gap, 3 × 3 mm in-plane resolution, and with TR of 2500 ms, TE of 35 ms, voxel size of 3 × 3 x 3 mm, flip angles of 90 degrees, as well as field of view of 240 mm^2^.

#### fMRI data preprocessing

The fMRI data from the emotional reactivity task was analyzed utilizing SPM8 (Wellcome Department of Cognitive Neurology, London, England) in conjunction with a preprocessing and analysis toolbox (http://github.com/ddwagner/SPM8w). The first step in the analysis involved correcting for differences in acquisition time between slices. Subsequently, within the first functional run, image realignment was performed using affine registration with 6 DOFs for motion correction. Following motion correction, the images underwent an unwarping procedure to decrease residual motion-related distortions that were not previously corrected. The next step involved normalizing the images to a standard stereotaxic space (3 × 3 x 3 mm isotropic voxels), SPM8 EPI template, which conforms to the ICBM 152 brain template space. Lastly, spatial smoothing was applied using a 6-mm full-width-at-half-maximum Gaussian kernel.

For each participant, a general linear model was computed, incorporating condition effects and covariates of no interest (a session mean, a linear trend to account for low-frequency drift, and six movement parameters derived from realignment corrections). The model was then convolved with a canonical hemodynamic response function. Subsequently, beta images were generated for the positive, negative, and neutral conditions, which were further used for the following analyses. To investigate potential associations between individual variations in state anxiety and corresponding variations in neural representations for each emotion condition, we used a whole-brain parcellation with 50 non-overlapping regions-of-interest (ROIs), which has been used in previous studies (14, 24). These 50 non-overlapping ROIs were defined based on functional coactivations from the Neurosynth database (25), and these ROI parcellations can be accessed at (https://neurovault.org/collections/2099/). The neural representations from beta maps for each valence condition were then extracted from each ROI. All subsequent analyses were performed using these neural representations obtained from these ROIs, with particular focus given to results from the fusiform gyrus, amygdala, vmPFC, and dmPFC. For completeness, exploratory analyses were performed on all other ROIs.

#### Intersubject representational similarity analysis (ISRSA)

We then employed ISRSA to explore which brain regions may reveal significant associations between individual variations in state anxiety and variations in neural representations for each valence condition by using the Nltools package (26). ISRSA investigates how intersubject variability in self-reported measurements relates to variability in neural representations. ISRSA accomplishes this by utilizing second-order statistics similar to classical representational similarity analysis (27, 28).

The ISRSA procedure begins with computing subject-by-subject distance matrices based on self-reported data (i.e., state anxiety or trait anxiety) and neural data, respectively. Regarding individual variability in state anxiety, in order to test whether compared to taking a single aggregate score, using a multidimensional representation would be better to capture individual variations in state anxiety, we conducted both multivariate distance and univariate aggregate score distance matrices for the state anxiety scores. For the multivariate distance matrix, cosine distance was utilized to calculate dissimilarity across all items in the state anxiety subscale between each pair of participants (29, 30). To obtain the univariate aggregated score distance matrix, we calculated the absolute difference between the composite state anxiety scores, thereby quantifying intersubject dissimilarity between each pair of participants. Regarding the neural data, within each valence condition, pairwise Euclidean distances between participants’ neural representations were computed per ROI, producing subject-by-subject distance matrices for each ROI. Spearman rank correlations were used to evaluate the associations between the intersubject dissimilarity of the behavioral multivariate and neural intersubject distance matrices, as well as the behavioral univariate aggregate score distance and neural intersubject distance matrices for each valence condition. To assess the significance of these correlations, Mantel permutation tests were utilized (28, 31), which is commonly employed to calculate one-tailed *p* values in ISRSA research (14, 32). The Mantel permutation test involves randomly reshuffling the order of participants in one multivariate distance matrix, recalculating the distance matrix, and then computing the correlation between the newly generated matrix and the original matrix (33). This process was repeated 10,000 times to generate a null distribution for correlation values. Based on the alternative hypothesis that the correlation value is greater than zero, the null distribution was used to calculate the *p*-value, allowing us to assess the significance of each correlation value using ISRSA. Given the increased probability of Type I error with multiple tests, Bonferroni correction was applied to control the family-wise error rate (FWE) during multiple comparisons. Consequently, the thresholded *p*-value after the Bonferroni correction was set to 0.001.

## Results

ISRSA results using multivariate distances of state anxiety consistently revealed significant associations between intersubject variations in state anxiety and neural representations in the fusiform gyrus. This effect was persistent across valence conditions: positive, *r*(1376) = 0.33, *p* < 0.001 (Figure 1(A-1)), negative (*r*(1376) = 0.32, *p* < 0.001; Figure 1(A-2)), and neutral scenes (*r*(1376) = 0.36, *p* < 0.001; Figure 1(A-3); Table 1). Similarly, significant associations were observed in area MT for positive (*r*(1376) = 0.36, *p* < 0.001; Figure 2(A)), negative (*r*(1376) = 0.32, *p* < 0.001; Figure 2(B)), and neutral scenes (*r*(1376) = 0.36, *p* < 0.001; Figure 2(C); Table 1). In addition, a significant association was observed in the insula for positive (*r*(1376) = 0.31, *p* < 0.001), but not in negative (*r*(1376) = 0.10, *p* = 0.094) nor neutral (*r*(1376) = 0.22, *p* = 0.004) scenes. Notably, a trending but non-significant association was observed in the amygdala during the viewing of negative scenes (*r*(1376) = 0.20, *p* = 0.011, Table 1). However, no significant associations were found in the vmPFC or dmPFC during the viewing of positive, negative, or neutral scenes (Table 1). On the other hand, results from the ISRSA using univariate aggregate score distances of state anxiety indicated no significant associations in the fusiform gyrus for positive (*r*(1376) =-0.01, *p* = 0.549; Figure 1(B-1)), negative (*r*(1376) = -0.05, *p* = 0.796; Figure 1(B-2)), and neutral conditions (*r*(1376) = -0.11, *p* = 0.959; Figure 1(B-3); Table 1). Similarly, no significant associations were found in area MT, amygdala, vmPFC, or dmPFC during the viewing of positive, negative, or neutral scenes (Table 1). In addition, we found trending correlations between intersubject similarity in trait anxiety and similarities of neural representations in the fusiform gyrus across all three valence conditions, but none of these effects would remain significant after applying Bonferroni correction for multiple comparisons (Table S1 and Figure S1).

**Table 1.** Summary of ISRSA results using multivariate distances and univariate aggregate score distances for computing intersubject similarity in state anxiety. The values indicate the Spearman correlation from the ISRSA analysis and the values within the parentheses correspond to the *p* values.

| ISRSA using multivariate distances |  |  |  |  |  |
| --- | --- | --- | --- | --- | --- |
|  | fusiform gyrus | area MT | amygdala | vmPFC | dmPFC |
| Positive | 0.33 (< 0.001) | 0.36 (< 0.001) | 0.09 (0.147) | 0.04 (0.284) | 0.08 (0.171) |
| Negative | 0.32 (< 0.001) | 0.32 (< 0.001) | 0.20 (0.011) | 0.01 (0.422) | 0.02 (0.410) |
| Neutral | 0.36 (< 0.001) | 0.36 (< 0.001) | 0.11 (0.108) | 0.11 (0.083) | -0.09 (0.869) |
| ISRSA using univariate aggregate score distances |  |  |  |  |  |
|  | fusiform gyrus | area MT | amygdala | vmPFC | dmPFC |
| Positive | -0.01 (0.549) | 0.02 (0.375) | 0.06 (0.205) | 0.01 (0.444) | 0.02 (0.371) |
| Negative | -0.05 (0.796) | 0.03 (0.335) | 0.04 (0.288) | 0.01 (0.411) | 0.04 (0.276) |
| Neutral | -0.11 (0.959) | -0.07 (0.882) | 0.11 (0.058) | 0.04 (0.234) | 0.06 (0.167) |

**Fig. 1.**
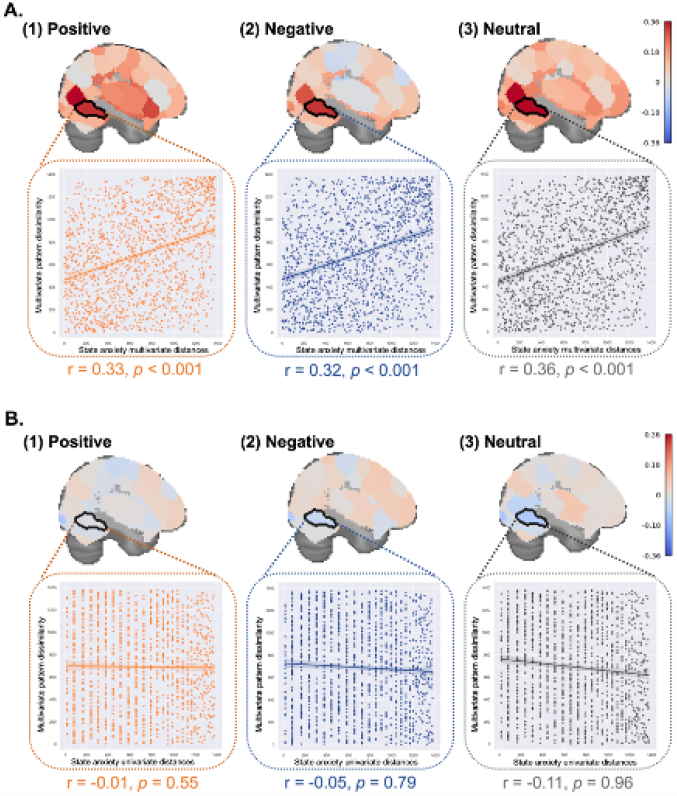
ISRSA using multivariate distances and univariate aggregate score distances in computing intersubject similarity in state anxiety. (A) ISRSA using multivariate distances revealed significant associations between individual variations in state anxiety and neural representations in the fusiform gyrus across all three valence conditions. (B) In contrast, ISRSA using univariate aggregate score distances yielded no significant associations in the fusiform gyrus. Spearman rank correlations and Mantel permutation tests were used to test for statistically significant brain-anxiety associations. The color bar indicates the Spearman correlation value from the ISRSA for each ROI.

**Fig. 2.**
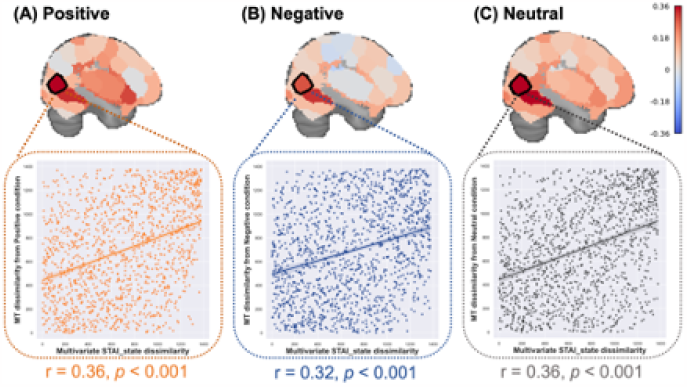
ISRSA using multivariate distances in computing intersubject similarity in neural representations from area MT. ISRSA using multivariate distances revealed significant associations between individual variations in state anxiety and neural representations in area MT across all three valence conditions.

## Discussion

In the present study, we demonstrate that intersubject similarity in neural representations of emotionally-charged and neutral stimuli was associated with similarity in their self-reported levels of state anxiety. Notably, these associations are primarily observed in the extrastriate cortex, specifically in the fusiform gyrus and area MT. Cognitive neuroscience research over the past two decades has emphasized the role of emotion in modulating stimulus representation in the visual cortex (19, 34, 35). By placing our results within this broader context, the present findings suggest that elevated state anxiety may influence perception by altering the neural representations of visual information in the extrastriate cortex (more so for the fusiform gyrus than the unexpected effect in the area MT). This proposition aligns with a recent study indicating that representations of valence are organized in a gradient within the fusiform gyrus (18). It is likely that an anxious state, compared to a calm state, may be associated with individual differences in perceiving emotionally charged stimuli differently (36, 37), which are related to differences in neural representations within the fusiform gyrus. Of note, our data show that state anxiety was significantly correlated with extrastriate cortex activity to both negative and positive scenes. These results describe the valence-general manner in which state anxiety might influence the neural representations of visual information, implying that even positively charged emotional scenes may be processed differently in the extrastriate cortex during an anxious state. This is in line with behavioral studies suggesting that nonclinical levels of anxiety may induce attentional bias for positive emotional visual stimuli (38, 39). Furthermore, this effect may not only be limited solely to emotionally-charged scenes but could extend to neutral scenes as well. Interestingly, this suggestion finds support in previous research demonstrating that differential neural and behavioral responses to emotionally neutral stimuli are associated with individual differences in anxiety (40, 41). It is also possible that individual differences in their levels of anxious states are associated with how low-level features are processed in the visual cortex. Overall, our study builds upon prior work that has focused on specific brain regions like the amygdala (12) and proposes a potential candidate mechanism for how anxious states might alter the representations of visual stimuli at the level of the extrastriate visual cortex.

Somewhat unexpectedly, there was a notable absence of significant associations in either the amygdala or vmPFC. While previous studies found correlations between state anxiety and neural responses using univariate methods, they usually involved facial expressions of emotion (10, 12) rather than emotional scenes. In cases where emotional scenes were used, such effects appear to depend on experimental design (passive viewing vs. cognitive reappraisal) or pre-existing mental health conditions (42). Our multivariate ISRSA approach expands on these findings to suggest that the extrastriate cortex is also impacted by state anxiety. That said, we do note that amygdala activity patterns to negative stimuli were marginally correlated with state anxiety (*p* < 0.011), but we are careful not to interpret this observation as this effect did not pass the corrected statistical threshold.

In sum, the present study examined the impact of a transient state of anxiety on neural representations of visual information. Rather than relying solely on a univariate aggregated score, we employed multivariate distances to capture individual differences in state anxiety. This approach considers the full spectrum of individual variations across all items, avoiding the reduction of the multivariate nature to a single average score. In line with other recent work (14), the present findings provide further evidence that, compared to a single aggregated score, the existence of more nuanced and sensitive individual differences might be within a multi-dimensional space (43). Lastly, although we include participants with diverse racial identities, the diversity in age distribution in our study is limited. Future studies are needed to expand the age distribution in order to test the generalizability of our findings. We are also aware that the sample size of neuroimaging studies is a crucial factor to be considered when conducting any analysis involving individual variations (44, 45). Although the sample size of the present study is comparable to recent studies utilizing the ISRSA approach to examine intersubject similarities between neural representations and other measurements (32, 46–50), the sample size should be carefully considered in future studies. In addition, a notable limitation of utilizing ISRSA should also be considered for future studies. As ISRSA only calculates the correlation between two distance matrices, this method contains no predictive capability like other predictive brain models derived based on machinelearning algorithms. Inferences drawn from the ISRSA are based on correlation rather than predictive power and causal relationships. Another limitation of our experimental design is that we only administered the questionnaire right after the scanning session, future studies employing an experimental design that causally manipulates state anxiety would be able to address this issue.

## Conclusion

These findings suggest a possible influence of state anxiety on extrastriate cortical activity, which in turn may shape how we perceive and interpret the world around us. Of course, these speculations should be formally tested in follow-up experimental work. Our data provide a useful starting point for future studies that might aim to experimentally induce anxious states and document the link between extrastriate cortical activity and the perception of emotional scenes.

## ACKNOWLEDGEMENTS

This research was supported by funding from the NSTC and MOE (MOST 109-2636-H-002-006 and 110-2636-H-002-004; NSTC 111-2628-H-002-004 and 111-2423-H-002-008-MY4; MOE NTU-CC-110L9A00702 and 112L9A00402 to P.-H.A. Chen) in Taiwan. This research was also supported by the National Research Foundation of Korea (NRF-2021R1F1A1045988 to M.J. Kim).

## Reference

1. Sonia J Bishop. Neurocognitive mechanisms of anxiety: an integrative account. Trends Cogn. Sci., 11(7):307–316, July 2007.

2. Andrew S Fox and Alexander J Shackman. The central extended amygdala in fear and anxiety: Closing the gap between mechanistic and neuroimaging research. Neurosci. Lett., 693:58–67, February 2019.

3. Dan W Grupe and Jack B Nitschke. Uncertainty and anticipation in anxiety: an integrated neurobiological and psychological perspective. Nat. Rev. Neurosci., 14(7):488–501, July 2013.

4. Catherine A Hartley and Elizabeth A Phelps. Changing fear: the neurocircuitry of emotion regulation. Neuropsychopharmacology, 35(1):136–146, January 2010.

5. M Davis and P J Whalen. The amygdala: vigilance and emotion. Mol. Psychiatry, 6(1): 13–34, January 2001.

6. Andrea M Cataldo and Andrew L Cohen. The effect of emotional state on visual detection: A signal detection analysis. Emotion, 15(6):846–853, December 2015.

7. Elizabeth A Phelps, Sam Ling, and Marisa Carrasco. Emotion facilitates perception and potentiates the perceptual benefits of attention. Psychol. Sci., 17(4):292–299, April 2006.

8. Amit Etkin and Tor D Wager. Functional neuroimaging of anxiety: a meta-analysis of emotional processing in PTSD, social anxiety disorder, and specific phobia. Am. J. Psychiatry, 164(10):1476–1488, October 2007.

9. M Justin Kim, Rebecca A Loucks, Amy L Palmer, Annemarie C Brown, Kimberly M Solomon, Mark Muraven, and Paul J Whalen. The structural and functional connectivity of the amygdala: From normal emotion to pathological anxiety. Behav. Brain Res., 223(2):403–410, October 2011.

10. Sonia J Bishop, John Duncan, and Andrew D Lawrence. State anxiety modulation of the amygdala response to unattended threat-related stimuli. J. Neurosci., 24(46):10364–10368, November 2004.

11. Jong Moon Choi, Srikanth Padmala, and Luiz Pessoa. Impact of state anxiety on the interaction between threat monitoring and cognition. Neuroimage, 59(2):1912–1923, January 2012.

12. Leah H Somerville, Hackjin Kim, Tom Johnstone, Andrew L Alexander, and Paul J Whalen. Human amygdala responses during presentation of happy and neutral faces: correlations with state anxiety. Biol. Psychiatry, 55(9):897–903, May 2004.

13. Caterina Gratton, Timothy O Laumann, Ashley N Nielsen, Deanna J Greene, Evan M Gordon, Adrian W Gilmore, Steven M Nelson, Rebecca S Coalson, Abraham Z Snyder, Bradley L Schlaggar, Nico U F Dosenbach, and Steven E Petersen. Functional brain networks are dominated by stable group and individual factors, not cognitive or daily variation. Neuron, 98(2):439–452.e5, April 2018.

14. Pin-Hao A Chen, Eshin Jolly, Jin Hyun Cheong, and Luke J Chang. Intersubject representational similarity analysis reveals individual variations in affective experience when watching erotic movies. Neuroimage, 216:116851, August 2020.

15. Pin-Hao Andy Chen and Yang Qu. Taking a computational cultural neuroscience approach to study parent-child similarities in diverse cultural contexts. Frontiers in Human Neuroscience, 15:703999, 2021.

16. Emily S Finn, Enrico Glerean, Arman Y Khojandi, Dylan Nielson, Peter J Molfese, Daniel A Handwerker, and Peter A Bandettini. Idiosynchrony: From shared responses to individual differences during naturalistic neuroimaging. Neuroimage, 215:116828, July 2020.

17. Wonyoung Kim and M Justin Kim. Morphological similarity of amygdala-ventral prefrontal pathways represents trait anxiety in younger and older adults. Proc. Natl. Acad. Sci. U. S. A., 119(42):e2205162119, October 2022.

18. Alison M Mattek, Daisy A Burr, Jin Shin, Cady L Whicker, and M Justin Kim. Identifying the representational structure of affect using fMRI. Affect Sci, 1(1):42–56, March 2020.

19. J S Morris, K J Friston, C Büchel, C D Frith, A W Young, A J Calder, and R J Dolan. A neuromodulatory role for the human amygdala in processing emotional facial expressions. Brain, 121 (Pt 1):47–57, January 1998.

20. Patrik Vuilleumier and Gilles Pourtois. Distributed and interactive brain mechanisms during emotion face perception: evidence from functional neuroimaging. Neuropsychologia, 45(1): 174–194, January 2007.

21. D D Wagner and T F Heatherton. Self-regulatory depletion increases emotional reactivity in the amygdala. Soc. Cogn. Affect. Neurosci., 8:410–417, July 2012.

22. Peter J Lang, Margaret M Bradley, Bruce N Cuthbert Cuthbert, Mark Greenwald, Arne Öhman, Dieter Vaitl, Alfons Hamm, Ed Cook, Andy Bertron, Margaret Petry, Rob Bruner, Mark Mcmanis, Deana Zabaldo, Shelley Martinez, Scott Cuthbert, Debbie Ray, Kathy Koller, Misty Kolchakian, Mary Pappenheimer, Alan Calpe, Sarah Eichler, Sarah Hayden, Marie Karlsson, and Katie Barber. International affective picture system (IAPS):. https://gitlab.pavlovia.org/rsaitov/experimental-psycholoy-ltu-final/raw/d3b3ec5364d25179c983d82c944baf04e06fd7ee/IAPS.TechManual.1-20.2008.pdf. Accessed: 2023-4-21.

23. C D Spielberger, F Gonzalez-Reigosa, and others. The state-trait anxiety inventory. Rev. Ordem Med., 1971.

24. Luke J Chang, Eshin Jolly, Jin Hyun Cheong, Kristina M Rapuano, Nathan Greenstein, Pin-Hao A Chen, and Jeremy R Manning. Endogenous variation in ventromedial prefrontal cortex state dynamics during naturalistic viewing reflects affective experience. Science Advances, 7(17):eabf7129, April 2021.

25. Alejandro de la Vega, Luke J Chang, Marie T Banich, Tor D Wager, and Tal Yarkoni. Large-Scale Meta-Analysis of human medial frontal cortex reveals tripartite functional organization. J. Neurosci., 36(24):6553–6562, June 2016.

26. Luke Chang, Eshin Jolly, Jin Hyun Cheong, Anton Burnashev, and Andy Chen. cosan-lab/nltools: 0.3.11, December 2018.

27. N Kriegeskorte, M Mur, and P Bandettini. Representational similarity analysis–connecting the branches of systems neuroscience. Frontiers in systems …, January 2008.

28. Lauri Nummenmaa, Enrico Glerean, Mikko Viinikainen, Iiro P Jääskeläinen, Riitta Hari, and Mikko Sams. Emotions promote social interaction by synchronizing brain activity across individuals. Proc. Natl. Acad. Sci. U. S. A., 109(24):9599–9604, June 2012.

29. Huchang Liao and Zeshui Xu. Approaches to manage hesitant fuzzy linguistic information based on the cosine distance and similarity measures for HFLTSs and their application in qualitative decision making. Expert Syst. Appl., 42(12):5328–5336, July 2015.

30. Devira Anggi Maharani, Carmadi Machbub, Pranoto Hidaya Rusmin, and Lenni Yulianti. Improving the capability of Real-Time face masked recognition using cosine distance. In 2020 6th International Conference on Interactive Digital Media (ICIDM), pages 1–6, December 2020.

31. N Mantel. The detection of disease clustering and a generalized regression approach. Cancer Res., 27(2):209–220, February 1967.

32. Jeroen M van Baar, Luke J Chang, and Alan G Sanfey. The computational and neural substrates of moral strategies in social decision-making. Nat. Commun., 10(1):1483, April 2019.

33. Gang Chen, Yong-Wook Shin, Paul A Taylor, Daniel R Glen, Richard C Reynolds, Robert B Israel, and Robert W Cox. Untangling the relatedness among correlations, part i: Nonparametric approaches to inter-subject correlation analysis at the group level. Neuroimage, 142: 248–259, November 2016.

34. Luis Carretié, Uxía Fernández-Folgueiras, Fátima Álvarez, Germán A Cipriani, Manuel Tapia, and Dominique Kessel. Fast unconscious processing of emotional stimuli in early stages of the visual cortex. Cereb. Cortex, 32(19):4331–4344, September 2022.

35. Philip A Kragel, Marianne C Reddan, Kevin S LaBar, and Tor D Wager. Emotion schemas are embedded in the human visual system. Sci Adv, 5(7):eaaw4358, July 2019.

36. Colette R Hirsch, Frances Meeten, Charlotte Krahé, and Clare Reeder. Resolving ambiguity in emotional disorders: The nature and role of interpretation biases. Annu. Rev. Clin. Psychol., 12:281–305, 2016.

37. A Mathews and C MacLeod. Cognitive approaches to emotion and emotional disorders. Annu. Rev. Psychol., 45:25–50, 1994.

38. K Mogg and B Marden. Processing of emotional information in anxious subjects. Br. J. Clin. Psychol., 29(2):227–229, May 1990.

39. Bradley C Riemann and Richard J McNally. Cognitive processing of personally relevant information. Cognition and Emotion, 9(4):325–340, July 1995.

40. Rebecca E Cooney, Lauren Y Atlas, Jutta Joormann, Fanny Eugène, and Ian H Gotlib. Amygdala activation in the processing of neutral faces in social anxiety disorder: is neutral really neutral? Psychiatry Res., 148(1):55–59, November 2006.

41. K Lira Yoon and Richard E Zinbarg. Interpreting neutral faces as threatening is a default mode for socially anxious individuals. J. Abnorm. Psychol., 117(3):680–685, August 2008.

42. Bryan T Denny, Jin Fan, Xun Liu, Kevin N Ochsner, Stephanie Guerreri, Sarah Jo Mayson, Liza Rimsky, Antonia McMaster, Antonia S New, Marianne Goodman, Larry J Siever, and Harold W Koenigsberg. Elevated amygdala activity during reappraisal anticipation predicts anxiety in avoidant personality disorder. J. Affect. Disord., 172:1–7, February 2015.

43. Eshin Jolly and Luke J Chang. The flatland fallacy: Moving beyond Low–Dimensional thinking. Top. Cogn. Sci., 11(2):433–454, April 2019.

44. Scott Marek, Brenden Tervo-Clemmens, Finnegan J Calabro, David F Montez, Benjamin P Kay, Alexander S Hatoum, Meghan Rose Donohue, William Foran, Ryland L Miller, Timothy J Hendrickson, Stephen M Malone, Sridhar Kandala, Eric Feczko, Oscar Miranda-Dominguez, Alice M Graham, Eric A Earl, Anders J Perrone, Michaela Cordova, Olivia Doyle, Lucille A Moore, Gregory M Conan, Johnny Uriarte, Kathy Snider, Benjamin J Lynch, James C Wilgenbusch, Thomas Pengo, Angela Tam, Jianzhong Chen, Dillan J Newbold, Annie Zheng, Nicole A Seider, Andrew N Van, Athanasia Metoki, Roselyne J Chauvin, Timothy O Laumann, Deanna J Greene, Steven E Petersen, Hugh Garavan, Wesley K Thompson, Thomas E Nichols, B T Thomas Yeo, Deanna M Barch, Beatriz Luna, Damien A Fair, and Nico U F Dosenbach. Reproducible brain-wide association studies require thousands of individuals. Nature, March 2022.

45. Tamas Spisak, Ulrike Bingel, and Tor D Wager. Multivariate BWAS can be replicable with moderate sample sizes. Nature, 615(7951):E4–E7, March 2023.

46. Noa Katabi, Hadas Simon, Sharon Yakim, Inbal Ravreby, Tal Ohad, and Yaara Yeshurun. Deeper than you think: Partisanship-Dependent brain responses in early sensory and motor brain regions. J. Neurosci., 43(6):1027–1037, February 2023.

47. Kun Il Kim, Wi Hoon Jung, Choong-Wan Woo, and Hackjin Kim. Neural signatures of individual variability in context-dependent perception of ambiguous facial expression. Neuroimage, 258:119355, September 2022.

48. Mai Nguyen, Tamara Vanderwal, and Uri Hasson. Shared understanding of narratives is correlated with shared neural responses. Neuroimage, 184:161–170, January 2019.

49. Shawn A Rhoads, Elise M Cardinale, Katherine O’Connell, Amy L Palmer, John W VanMeter, and Abigail A Marsh. Mapping neural activity patterns to contextualized fearful facial expressions onto callous-unemotional (CU) traits: intersubject representational similarity analysis reveals less variation among high-CU adolescents. Personal Neurosci, 3:e12, November 2020.

50. Qiang Xu, Jiali Hu, Yi Qin, Guojie Li, Xukai Zhang, and Peng Li. Intention affects fairness processing: Evidence from behavior and representational similarity analysis of event-related potential signals. Hum. Brain Mapp., 44(6):2451–2464, April 2023.

